# G-Quadruplexes Act as Sequence Dependent Chaperones via Protein Oligomerization

**DOI:** 10.1101/850263

**Authors:** Adam Begeman, Theodore J. Litberg, Jennifer Bourne, Zhenyu Xuan, Scott Horowitz

**Affiliations:** Department of Chemistry & Biochemistry, Knoebel Institute for Healthy Aging, University of Denver, Denver, Colorado 80208, United States; Department of Cell and Developmental Biology, University of Colorado School of Medicine, Aurora, Colorado 80045 United States; Department of Biological Sciences, Center for Systems Biology, University of Texas at Dallas, Richardson, TX 75080, United States

## Abstract

Maintaining proteome health is important for cell survival. Nucleic acids possess the ability to prevent aggregation up to 300-fold more efficiently than traditional chaperone proteins. In this study, we explore the sequence specificity of the chaperone activity of nucleic acids. Evaluating over 500 nucleic acid sequences’ effects on aggregation, we demonstrate that the holdase chaperone effect of nucleic acids is highly sequence dependent. Quadruplexes are found to have especially potent effects on aggregation with many different proteins via quadruplex:protein oligomerization. These observations contextualize recent reports of quadruplexes playing important roles in aggregation-related diseases, such as Fragile X and Amyotrophic lateral sclerosis (ALS).

## Main Text

Chaperones are a diverse group of proteins and other molecules that regulate proteostasis (1) in the cell by preventing protein aggregation (holdases) and helping protein folding (foldases). Recently, molecules other than traditional protein chaperones have been shown to play important roles in these processes (2, 3). We recently showed that nucleic acids can possess potent holdase activity, with the best sequences having higher holdase activity than any previously characterized chaperone (4). Nucleic acids can also collaborate with Hsp70 to help protein folding, acting similarly to small heat shock proteins (4–7). Nucleic acids can also bring misfolded proteins to stress granules (8), and are a primary component of the nucleolus, which was recently shown to store misfolded proteins under stress conditions (9). However, the structural characteristics, sequence dependence, and mechanistic understanding of how nucleic acids act as chaperones remains unclear.

A critical question in understanding the holdase activity of nucleic acids is whether this activity is sequence specific? Previously, we showed that polyA, polyT, polyG, and polyC prevented aggregation with varying kinetics, suggesting that sequence specificity was possible (4). Here, we test sequence specificity by examining over 500 nucleic acids of varying sequence for holdase activity. The holdase activity is found to be highly sequence specific, with quadruplexes showing the greatest activity. Several quadruplexes displayed generality, with potent holdase activity for a variety of different proteins. Further examination of these quadruplex sequences demonstrated that the holdase activity largely arose through quadruplex:protein oligomerization. These results help explain several recent reports of quadruplex sequences playing important roles in oligomerization, aggregation, and phase separation in biology and pathology, and that these are common properties of quadruplex interactions with partially unfolded or disordered proteins.

### Sequence Specificity of Holdase Activity

To determine the sequence specificity of the holdase activity of nucleic acids, we measured light scattering and turbidity via absorbance in a thermal aggregation assay (Fig. 1A) for 312 nucleic acid sequences (Fig. 1B). These nucleic acids were nearly all 20 bases in length, single stranded DNA (ssDNA) sequences of random composition. Bulk DNA was used as a positive control (4). Plotting the percent aggregation for each sequence demonstrates that the holdase activity of the ssDNA is very sequence dependent (Fig. 1B). Sequences nearly spanned the complete range in activity, from barely affecting aggregation, to nearly preventing all protein aggregation for over an hour.

**Fig. 1.**
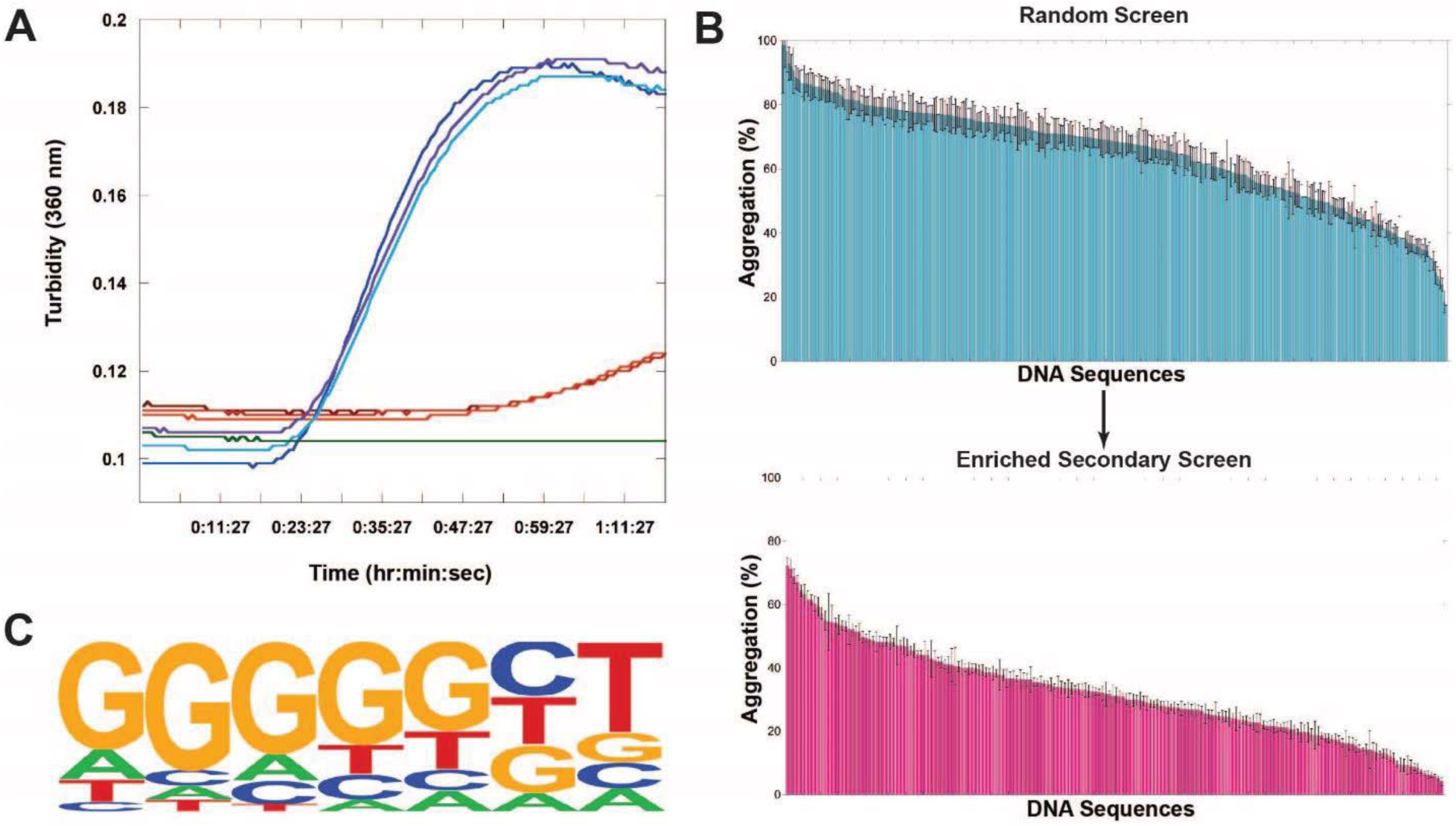
(A) Representative example citrate synthase protein aggregation assay. Turbidity and light scattering were measured in a multimode plate reader at 360 nm for 1.5 hours of incubation at 50° C. Blue lines represent triplicate citrate synthase alone, red and orange lines represent triplicate citrate synthase incubated with a single ssDNA sequence, green is buffer alone. (B) Screen of ssDNA sequences for holdase chaperone activity. Each bar represents a different 20-nt sequence, sorted by activity. Aggregation % was measured as the normalized average of triplicate citrate synthase turbidity measurements after 1.5 hours of incubation at 50°C. Lower aggregation indicates greater holdase function. The initial screen used random, non-redundant sequences (top), which led to a followup enriched screen (bottom). Error bars are SE. (C) HOMER Logo of motif found by analyzing screen (statistics: p<1.0×10^-13^, FDR<0.001, % of Targets: 53.85, % of Background: 7.69).

With this high level of sequence dependence, we next performed bioinformatics to determine if any sequence motifs encoded holdase activity. We first found that the holdase activity is positively correlated with the guanine content in the sequences (ρ=0.24, p-value=1.5×10^-5^). Comparing the top third in holdase activity to the bottom third, only one motif was found to be significantly enriched in sequences with higher holdase activity (53.85% vs 7.69%, FDR =0.001) (Fig. 1C). This motif contains five consecutive guanines followed by any base and then thymine. A similar G-rich motif (consensus pattern: BGGSTGAT) was also found by a regression based method (R2 = 0.61, p-value =1×10^-5^).This analysis suggested that the most potent holdase activity was encoded by a polyG motif.

To verify this polyG motif, we tested another 192 sequences for holdase activity that had high guanine content. These sequences include 96 sequences with a 55% bias towards guanine bases, 40 sequences with a 75% bias towards guanine bases, and 56 having different positional variations of the aforementioned polyG motif (Fig. 1C). Comparing the original random sequences to these guanine-rich sequences, the average aggregation was substantially reduced in the enriched guanine set, from 64.8% to 32.0%. Within the enriched guanine set, however, there was still a great deal of variation, with the data spanning aggregation from 72% to 4%. This wide variability suggested that the motif required more than just high guanine content. Within the subset of sequences with a 55% bias towards guanine, a significant polyG motif was again identified by comparing sequences having different holdase activity. Within the subset of 75% guanine-containing sequences, no statistically significant differences were found, as most sequences contained at least one polyG motif. We also tested holdase activity for 56 sequences having different positional variations of the aforementioned polyG motif, which did not find positional dependency within the sequence for the holdase activity. In the best sequences from this enriched assay, the nucleic acid completely prevented protein aggregation for the entire hour and a half experiment. These results confirmed that the holdase activity was associated with a polyG motif.

### G-Quadruplexes as Potent Holdases

PolyG is well known to form quadruplexes when provided with appropriate counter ions. Composed of polyG bases forming pi-stacked tetrads, guanine quadruplexes are a class of structured nucleic acids that have been of increasing interest due to their regulatory role in replication, transcription, and translation (10, 11). Quadruplexes have also recently been implicated in several protein aggregation genetic disorders, such as Fragile X syndrome and ALS (12–17).

To test if the sequences containing polyG that had potent holdase activity were forming quadruplexes, we performed circular dichroism (CD) spectroscopy experiments on three sequences to determine their secondary structure. The CD spectra showed distinct peaks at 260 and 210 nm, with a trough at 245 nm, indicative of parallel quadruplex formation (Fig. 2A) (18), and distinct from a control sequence (sequence 42) that had poor chaperone activity and no polyG motif. This supposition was further supported by examining the emission spectra of N-methylmesoporphyrin IX (NMM), a well characterized parallel quadruplex binding fluorophore (19, 20). The NMM spectra indicated that all three sequences formed parallel quadruplexes at the concentration used in aggregation assays, unlike the ssDNA control (Fig. 2B). Melting experiments indicated that these quadruplex structures were stable, with 91% of the quadruplex structure remaining at 50° C, the temperature used in the holdase assays. These experiments confirmed that the holdase activity of these polyG-containing sequences were associated with quadruplex structure. Re-analyzing the heat denaturation aggregation assay data presented above, of the 160 sequences tested that had the sequence properties to form quadruplexes, 133 appeared in the top third of data, making up 79% of the sequences in that subset. 152 of the 160 quadruplex sequences also decreased aggregation by at least 50%.

**Fig. 2.**
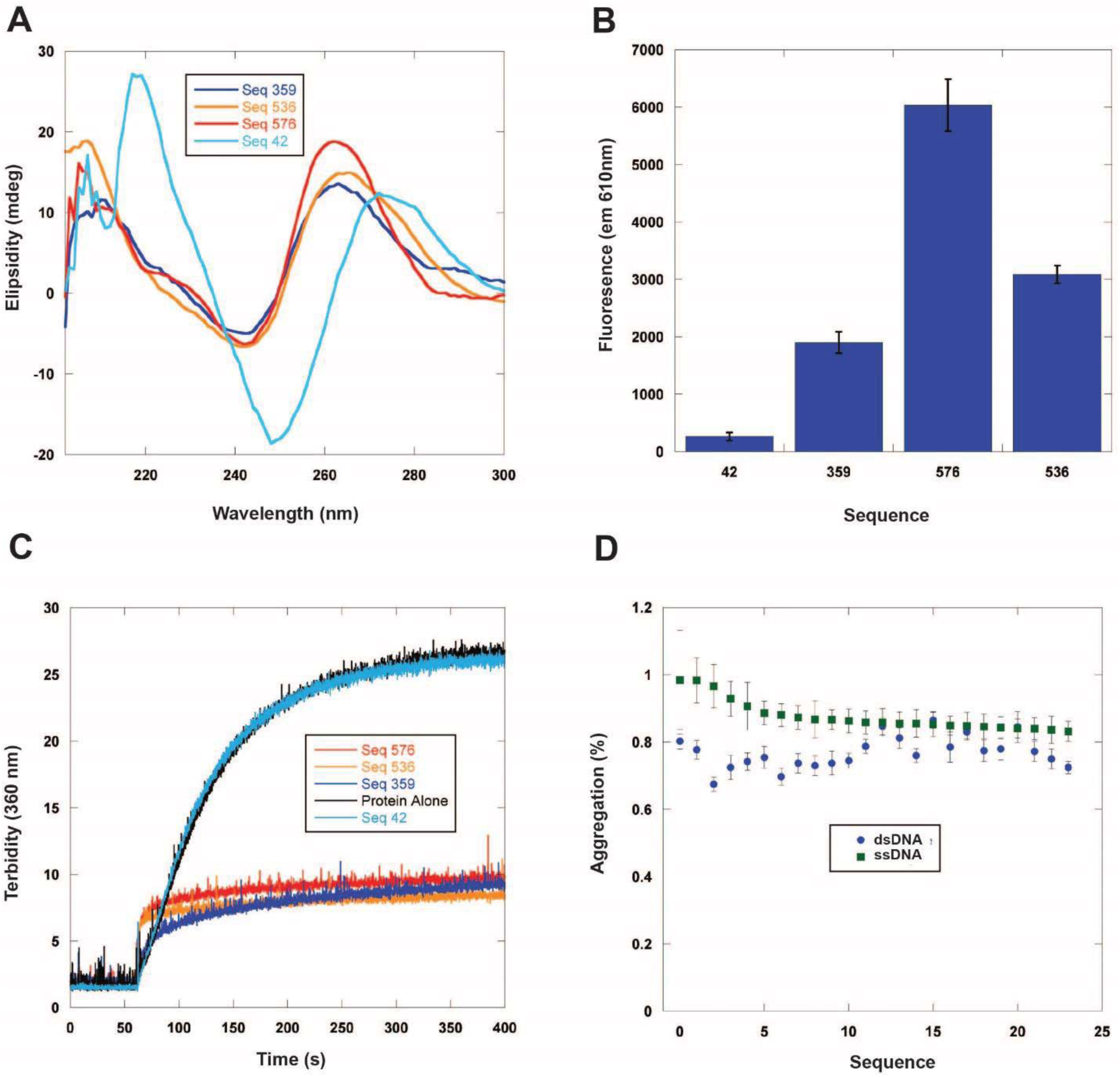
Characterization of quadruplex content and holdase activity. Sequences 359, 536, and 576 all displayed holdase activity and contain a polyG motif. Sequence 42 was used as a negative control, as it performed poorly as a holdase chaperone and did not contain a polyG motif. (A) Structural characterization of holdase nucleic acids using circular dichroism. Peaks are observed at 260 nm and 210 nm, as well as a trough at 245 nm, indicating the presence of parallel G-quadruplexes. Thermal stability of quadruplexes shown in Fig. S2. (B) NMM fluorescence measured at 610 nm. (C) Aggregation during chemically induced aggregation via right angle light scattering at 360 nm. Concentration dependence for Seq 359 shown in Fig. S3. (D) Holdase activity of ssDNA compared to its duplexed counterpart in an identical aggregation assay to that shown in Fig. 1.

We further characterized the holdase activity of the quadruplex-forming sequences using chemical denaturation aggregation assays in which the protein starts in a denatured state. Light scattering experiments confirmed the holdase activity in at least nine different quadruplex sequences (Fig. 2C, Fig. S1). This data also suggests that the quadruplexes are binding a misfolded or partially denatured form of the protein rather than the native state.

The higher level of activity from quadruplex DNA raised the question of whether any structured DNA could have a similar effect. In other words, could the activity arise from any DNA with greater structure than ssDNA? To test this possibility, we tested the holdase activity of 24 duplexed sequences to compare directly with their single-stranded counterparts. Overall, the differences were small, and in many cases statistically insignificant (Fig. 2D). These experiments suggest that the holdase activity displayed here could be specific to quadruplex structures, and not other structured DNAs.

### Generality of Holdase Activity

To determine the generality of this quadruplex holdase activity, we performed aggregation assays with three other proteins, luciferase, lactate dehydrogenase (LDH), and malate dehydrogenase (MDH). To check whether this holdase activity was quadruplex-specific with multiple proteins, we tested 16 quadruplex sequences and 8 single-stranded sequences with each protein. These proteins have varying structural properties, ranging in pI from 6.1 to 8.5, and size from 62.9 kD to 140 kD. With all four proteins, the quadruplexes severely decreased protein aggregation, demonstrating strong holdase activity. For luciferase, LDH, and MDH, most of the quadruplex sequences tested were able to completely prevent protein aggregation. However, the single-stranded sequences demonstrated little to no activity for all of these proteins (Fig. 3).

**Fig. 3.**
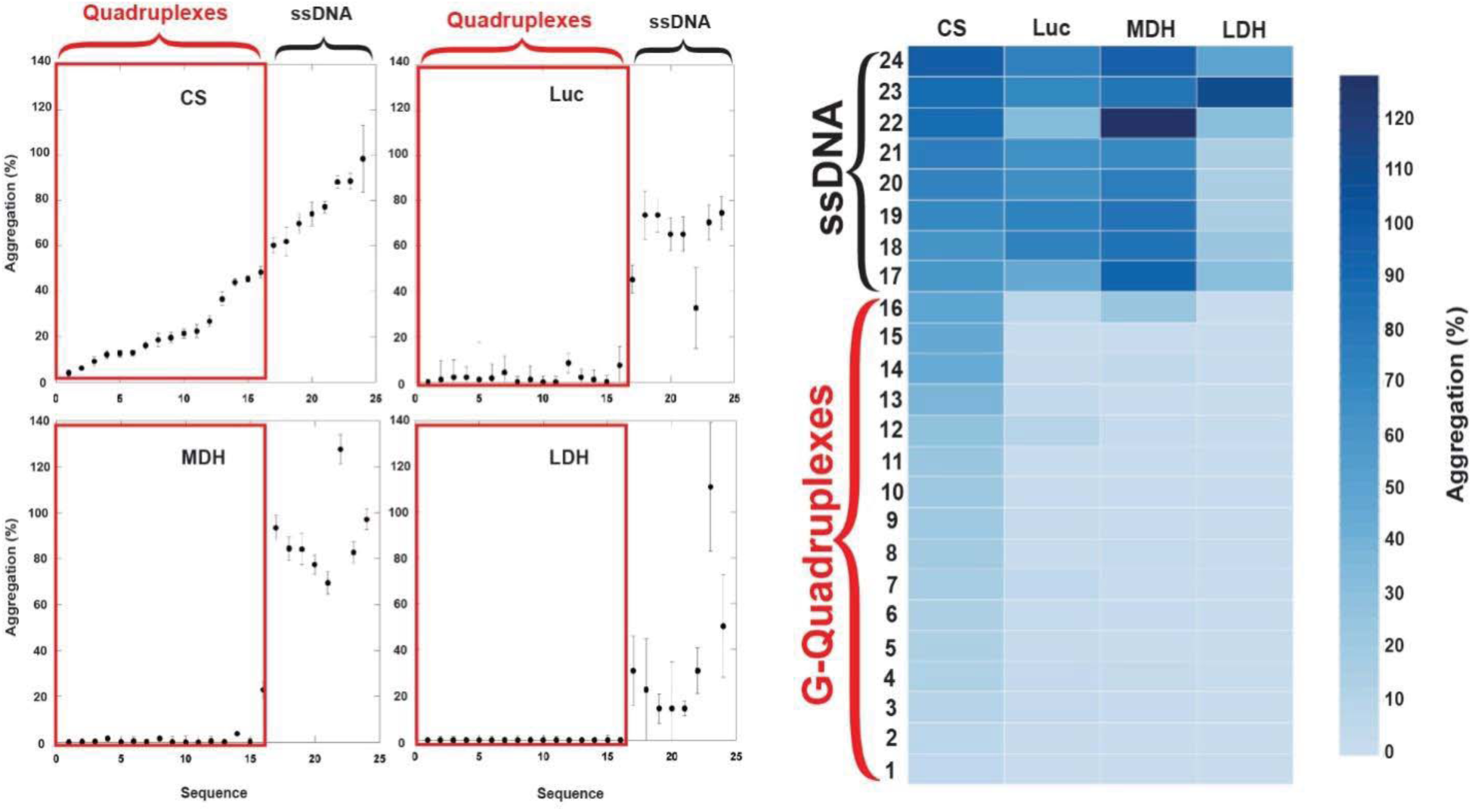
Generality of G-quadruplex holdase activity using four different proteins: Luciferase (Luc), Citrate Synthase (CS), L-Malate Dehydrogenase (MDH), and L-Lactate Dehydrogenase (LDH) Boxed in red are the 16 sequences with the propensity to form quadruplexes, while the remaining 8 sequences are non-structured ssDNA (left). This data is also shown as a heat map (right).

These data strongly suggest that the holdase activity displayed by quadruplex sequences is general, while also unique to quadruplex-forming sequences. Of note, LDH is the only of these proteins with previously characterized DNA-binding activity towards both duplex and ssDNA (21–23), but its aggregation was only significantly reduced by binding to quadruplex sequences (Fig. 3).

### Holdase Activity Due to Oligomerization

While analyzing the light scattering data from chemically denatured citrate synthase in the presence of quadruplexes, we noticed that although the total light scattering was greatly decreased by the presence of the quadruplexes, the quadruplexes caused a small initial jump in light scattering (Fig. 4A). These data are highly reminiscent of the pattern we observed recently in which nucleic acids could prevent protein aggregation by promoting protein:nucleic acid oligomerization (24).

**Fig. 4.**
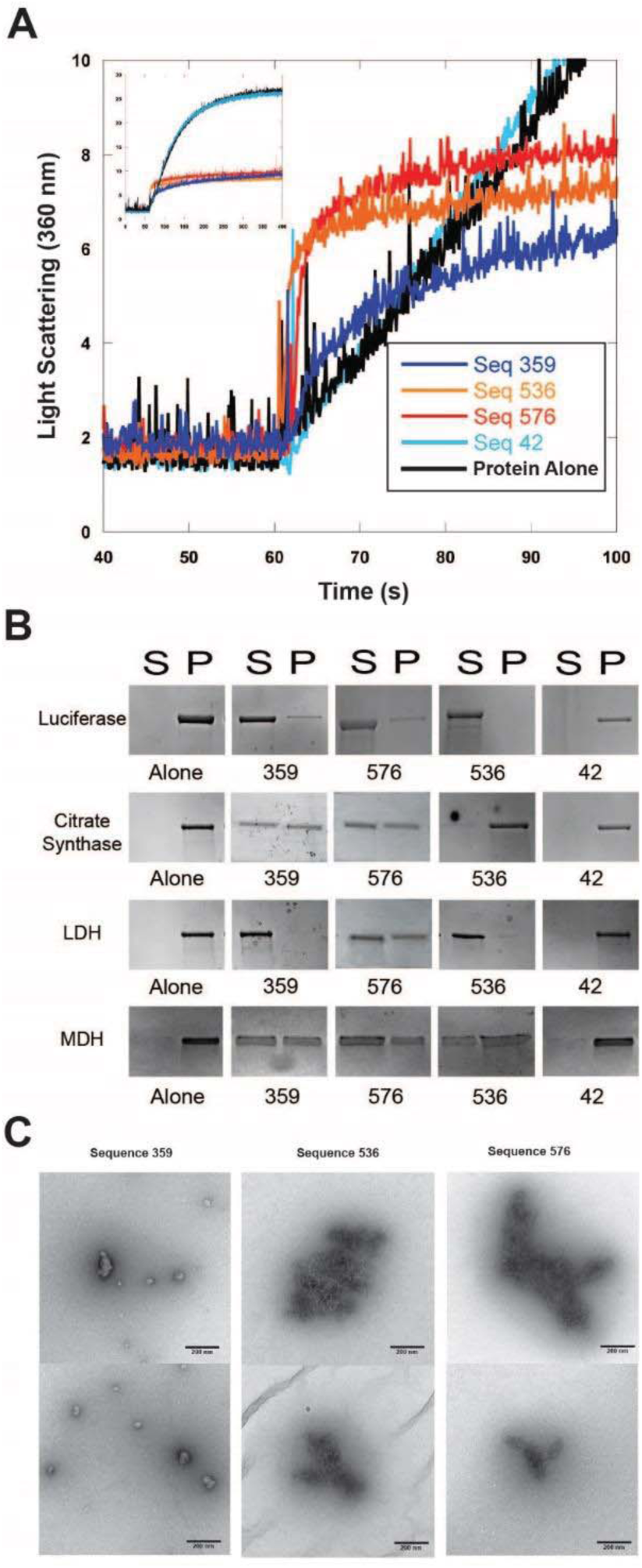
Quadruplex-containing sequences promote oligomerization. Sequences 359, 536, and 576 all displayed holdase activity and contain a polyG motif. Sequence 42 was used as a negative control, as it performed poorly as a holdase chaperone and did not contain a polyG motif. (A) Right angle light scattering of chemically induced aggregation of citrate synthase. Initial kinetics of aggregation shown with full time scale as the insert. (B) Spin down assay with four different proteins denatured at 60 °C in the presence of DNA (C) Transmission electron microscopy negative stain images of soluble fractions from chemically induced aggregation spin down assays. Citrate synthase oligomers were observed in each of the quadruplex cases, although the morphology of the quadruplex was dependent on the DNA sequence. Correlating chemical denaturation spin down assays are shown in Fig. S5.

To determine whether the quadruplexes tested here were acting in a similar manner, we performed additional spin-down assays, CD, and transmission electron microscopy (TEM) experiments. Spin-down assays were performed by heating citrate synthase, luciferase, MDH, or LDH with quadruplexes to 60°C for 15 min, and then returning them to room temperature. The sample was centrifuged to then separate soluble and pellet fractions. SDS-PAGE gels demonstrated that the quadruplexes kept the proteins soluble even at extreme temperatures (Fig. 4B), similar to previously characterized oligomerization cases (24). Of note, a control single stranded sequence of the same length (sequence 42) did not keep the proteins soluble under these conditions, even for the well-characterized DNA binding protein LDH (Fig. 4). Measuring CD spectra of luciferase protein as a function of temperature in the presence of quadruplexes showed that the protein maintained partial β-sheet structure as high as 80°C, and that this non-native structure was retained upon return to room temperature (Fig. S4), similar to previous oligomerization cases (24). Finally, negative stain TEM imaging showed that the quadruplexes caused the formation of protein oligomers (Fig. 4C). Notably, the morphology of the oligomers varied with different quadruplex structures, suggesting that the sequence or structural details of the quadruplex influence the structure of the subsequent oligomerization that occurs.

## Discussion

In this study, a systematic investigation of the holdase activity of nucleic acids demonstrated that this activity is sequence specific, and that quadruplex sequences display potent holdase activity. This holdase activity was demonstrated to be general, preventing the aggregation of multiple proteins that differed considerably in pI, size, and function. Further testing showed that this activity arose largely via protein:nucleic acid oligomerization. To our knowledge, this activity was also found to be more efficient than any previously characterized protein chaperone (4).

Quadruplex sequences have recently been implicated in aggregation and phase separation events in the cell that are associated with pathology. The quadruplex-forming GGGGCC repeat expansion in the c9orf72 gene is thought to be a frequent cause of both ALS and frontotemporal dementia (FTD) (12, 25). This quadruplex sequence is transcribed into sense and anti-sense RNA that have been shown to sequester numerous RNA binding proteins into toxic intranuclear foci, resulting in aggregation (26–30). GGGGCC quadruplexes forming foci with disordered proteins in the cell is highly consistent with the results presented here. FMRP protein has been shown to be one of the leading causes of the fragile X syndrome (17) as well as one of the leading causes of monogenetic forms of autism (31, 32). In addition to aggregating in disease (33, 34), FMRP is a known quadruplex-binding protein (16, 17). Future studies on whether these roles are related would be of significant interest. The results presented here suggest that these disease-relevant cases are not unique, as this behavior is a general property of quadruplex interaction with partially unfolded or disordered proteins under stress conditions.

Quadruplexes preventing protein aggregation by oligomerizing with their clients is reminiscent of the action by small heat shock proteins (sHsps). This class of chaperones forms large hetero-oligomer complexes with different clients, keeping these clients soluble and in an accessible state for later refolding by ATP-dependent chaperones (5, 7). The quadruplexes studied here also formed stable and soluble oligomer complexes with partially folded proteins, suggesting that the quadruplexes could use a molecular mechanism similar to those of the sHsps. Of note, the morphology of the oligomers varied with quadruplex sequence, further suggesting that altering quadruplex sequence could be a way to control the oligomerization of proteins under stress conditions.

## Acknowledgments

The authors would like to acknowledge E. Chapman, J. Yesselman, and S. Barbee for helpful conversation.

## Author contributions

S.H. conceptualized and administered the project; all authors designed experiments and analyzed data, A.B., T.L., J.B., and Z.X. performed investigations, S.H. and A.B. wrote the paper with assistance from all authors.

## Data and materials

All data is available in the main text or the supplementary materials.

## Materials and Methods

### Sourcing DNA

All DNA was ordered from Integrated DNA Technology using their standard desalting and purification procedures. For the heat aggregation plate reader assays, DNA was ordered lyophilized, and normalized to guaranteed molar weights by IDT. This DNA was then resuspended in the given buffer and pipetted directly into the plate wells after thorough pipette mixing. The duplex DNA was pre-annealed by IDT using their standard annealing protocol (https://www.idtdna.com/pages/education/decoded/article/annealing-oligonucleotides). For all other experiments, DNA was ordered lyophilized in tube form from IDT at the maximum yield achieved during synthesis. The DNA was then resuspended and thoroughly mixed with the given buffer to a known concentration.

### Thermal Aggregation

For the initial thermal aggregation assays, 312 single stranded sequences of random sequence, 24 of which varied in length from 15 to 20 bases long, while the rest were all 20 bases long, were incubated with 500 nM Citrate Synthase from porcine heart (Sigma-Aldrich C3260-5KU) in a 1:2 protein:DNA strand concentration. Aggregation was measured absorbance at 360 nm in a Biotek Powerwave multi-mode plate reader, with shaking and measurements every 36 seconds. In every assay, the plates were transferred from ice to a preheated 50° C plate reader, and the temperature was held constant throughout the entire experiment. Each plate was run for 1.5 hours in 40 mM Hepes, 7.5 pH (KOH) buffer. The sequences were run in triplicate. Percent aggregation was calculated as a function of the maximum absorbance value recorded in the hour and a half divided by the maximum protein alone absorbance value. Error bars shown are standard error propagated from both the triplicate protein alone and triplicate experimental measurement. As a control, herring testes DNA (Sigma) was also run on each plate to ensure consistency of data.

The enriched population of 192 G-rich sequences (length 20 bases) was performed by biasing 96 of the sequences toward guanine bases at a rate of 55%. 40 sequences were biased towards guanine bases by 75%, and the remaining 56 sequences were created with the motif GGGGGNT systematically placed throughout the sequence, with the remaining 13 bases chosen at random. This process was accomplished by altering the random bias in our random sequence generating software, which can be found at https://github.com/adambegeman/IDT_DNA_Generator.

We also tested aggregation using Quantilum Recombinant Luciferase (Promega), L-malate dehydrogenase (MDH) from pig heart (Sigma-Aldrich 10127914001), and L-Lactate Dehydrogenase (LDH) from rabbit muscle (Sigma-Aldrich 10127876001). We chose 24 sequences from the citrate synthase assay whose anti-aggregation ability spanned the entire range of our data. Of these 24, 16 had a propensity to form quadruplexes. The assay with these other three proteins was carried out identically to the citrate synthase thermal denaturation assay, with LDH being run for 3 hours due to its higher stability at 50° C. These assays were run in 1:2 protein:DNA strand ratios using 10 mM sodium phosphate, 7.5 pH buffer at protein concentrations of 500 nM Luciferase, 2 μM MDH, and 4 μM LDH.

### Motif analysis

Motif analysis was performed using the HOMER package (35) to compare the one third of sequences with the highest holdase activity to the one third of sequences with the lowest holdase activity. The parameters used were “-len 5,6,7,8,9,10 -norevopp -noconvert -nomask -mis 2 - basic -nogo -noredun -noweight -fdr 1000”. We also used MatrixREDUCE (36) to identify motifs correlated with holdase activity by regression method. Spearman correlation analysis was performed by R.

### Chemical Aggregation

Procedure was adapted from *Docter et al.* and *Gray et al.* (2, 4). For the chemically induced aggregation, 12 μM Citrate Synthase was denatured in 6 M guanidine-HCl, 40 mM HEPES, for approximately 16 h at 23° C, then diluted to 75 nM into 40 mM HEPES, pH 7.5 (KOH), with constant stirring at 23° C in the presence of 150 nM 20-mer ssDNA. The resulting aggregation was then measured via right angle light scattering at 360 nm in a fluorimeter with constant mixing. Results were consistently repeated on three separate days, with representative curves shown.

### N-methylmesoporphyrin IX (NMM) Fluorescence

NMM is a well characterized fluorophore that increases fluorescence when bound to parallel quadruplexes (19). The emission spectra of 10 μM NMM was measured using an excitation wave length of 399 nm, and an emission range of 550 to 750 nm in the presence of 1 μM DNA. Samples were run in triplicate at 25° C in a multimode plate reader. Reported values are taken at 610 nm, the emission maxima, as a function of increase in fluorescence compared to a NMM alone triplicate control.

### Transmission Electron Microscopy (TEM)

For the oligomer TEM samples, chemical denaturation spin down assays were run using citrate synthase. 46.4 μM Citrate synthase was denatured in 4.8 M guanidine-HCl, 40 mM HEPES buffer for 16 hours. The citrate synthase was then diluted to 1.5 μM in a 100 μL sample containing 3 μM of the target sequence that induced oligomerization. 15 minutes after injection, the sample was spun down at 16,100 x g for 20 min at 4° C. The soluble portion was pipetted off and transferred on ice for TEM analysis.

A positively charged copper mesh grid coated in formvar and carbon (Electron Microscopy Sciences) using the PELCO easiGlow Discharge system was used for each soluble sample. The charged copper grids had 5 μL of sample applied for 20 seconds and then lightly blotted off using a Whatman filter paper. The grids were then rinsed using 2 drops of MilliQ water, with filter paper bloating for each wash. Finally, the grids were then stained using two drops of a 0.75% uranyl formate solution. The first drop served as a quick wash, followed by 20 seconds of staining using the second drop. The grids were then blotted and allowed to dry. The TEM images were captured using a FEI Tecnai G2 Biotwin TEM at 80 kV with an AMT side-mount digital camera. In order to better visualize the intricacies of each oligomer, the images’ contrast and brightness was uniformly enhanced using Adobe Photoshop.

### Circular Dichroism

The DNA CD spectra were obtained using a Jasco J-1100 circular dichroism at 23° C. The spectra were captured using 25 μM concentrations of DNA in 10 mM sodium phosphate, 7.5 pH buffer. The CD measurements were taken from 300 nm to 190 nm at 1 nm intervals using a 1 nm/sec scanning speed. The shown spectra are a product of three accumulations using the same conditions.

Melting curves for the quadruplex sequences were obtained using the same methods as described above with spectra taken at 10° C intervals from 20° C to 80° C at a ramp rate of 5° C/ min. The luciferase protein denaturation CD spectra were captured in protein:DNA ratios of 1:2 using 3.2 μM luciferase. The CD spectra were captured at 10° C intervals from 15° C to 85° C using a ramp rate of 3° C/ min. The reverse spectra was also captured, but the sequences failed to show any ability to recover native luciferase structure. Spectra were captured from 260 nm to 190 nm at 1 nm intervals using a 1 nm/sec scanning speed over two accumulations. The concentrations were chosen such that the DNA concentration was below the observable sensitivity range of the instrument.

### Spin Down Aggregation Assays

100 μL of 3.2 μM protein and 6.4 μM ssDNA were thermally denatured together at 60°C for 15 min in 10 mM sodium phosphate, pH 7.5 buffer. The resulting solution was then centrifuged at 16,100 x g for 20 minutes at 4° C to separate the soluble and insoluble fractions. After centrifugation, the supernatant (approximately 97 μL) was removed and the pellet resuspended using 1 mM β-mercaptoethanol (Fisher Scientific) in 1x TG-SDS buffer (Bio Basic Inc) to the original sample volume of 100 μL. 10 μL of the soluble and pellet fractions were then run on a denaturing SDS-PAGE gel and visualized using Coomassie blue. Gels were reproduced on three separate days, with representative assays shown.

For the chemical denaturation spin down assay, 46.4 μM citrate synthase was denatured in 4.8 M guanidine-HCl, 40 mM HEPES buffer for 16 hours. The citrate synthase was then diluted to 2.5 μM in a 100 μL sample containing 5 μM of the target sequence that induced oligomerization. After 5 minutes, the samples soluble and pellet fractions were separated via the same methods as the thermal denaturation spin down assays. The gels were then run in triplicate identically to the thermal denaturation experiments, with representative gels shown.

---

**Fig. S1.**
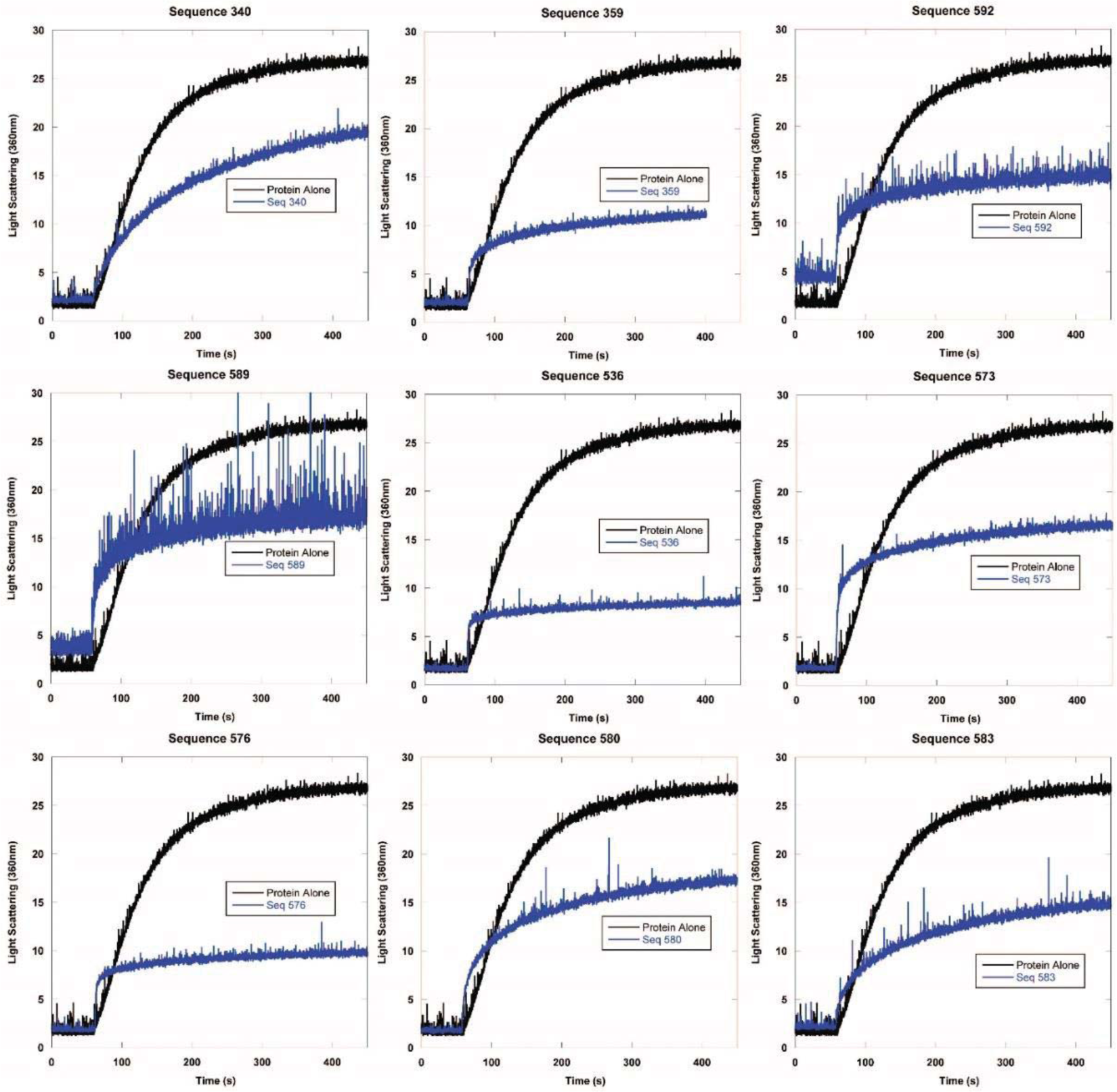
Representative examples of chemical aggregation tests of multiple quadruplex-containing sequences with citrate synthase. Concentration ratio is 1:2 protein to DNA strand.

**Fig. S2.**
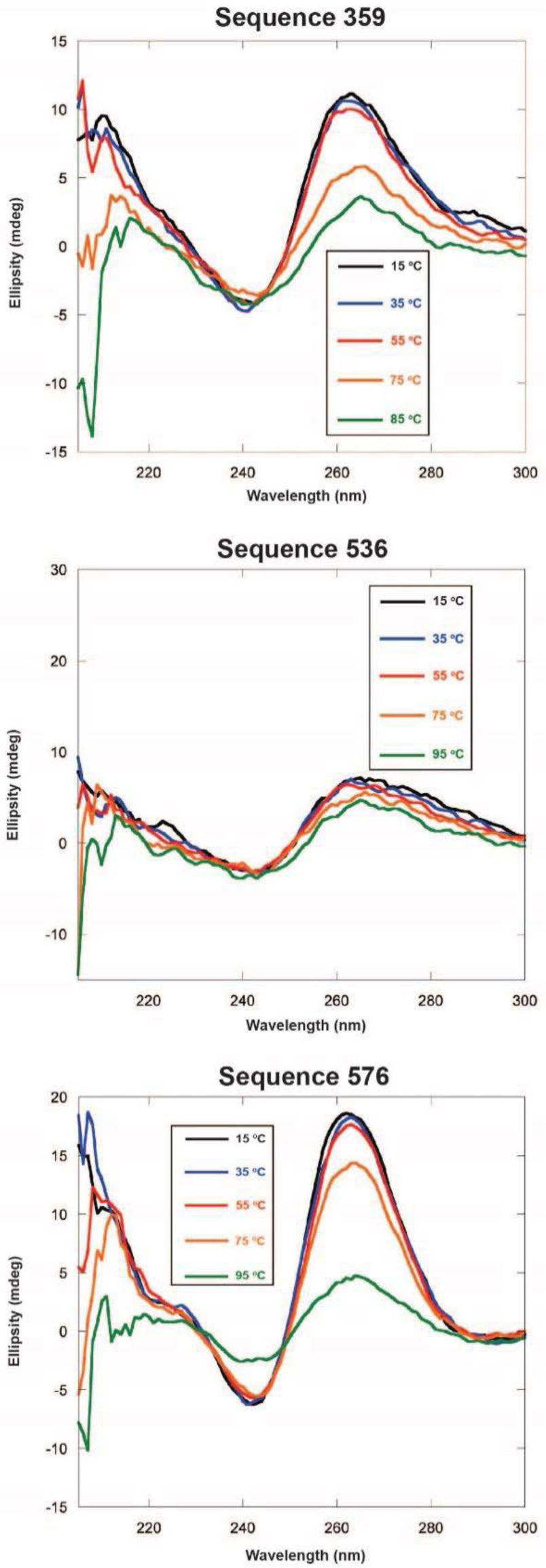
Thermal stability of quadruplex-containing sequences as measured by CD spectroscopy. Each line represents a wavelength scan at the indicated temperature.

**Fig. S3.**
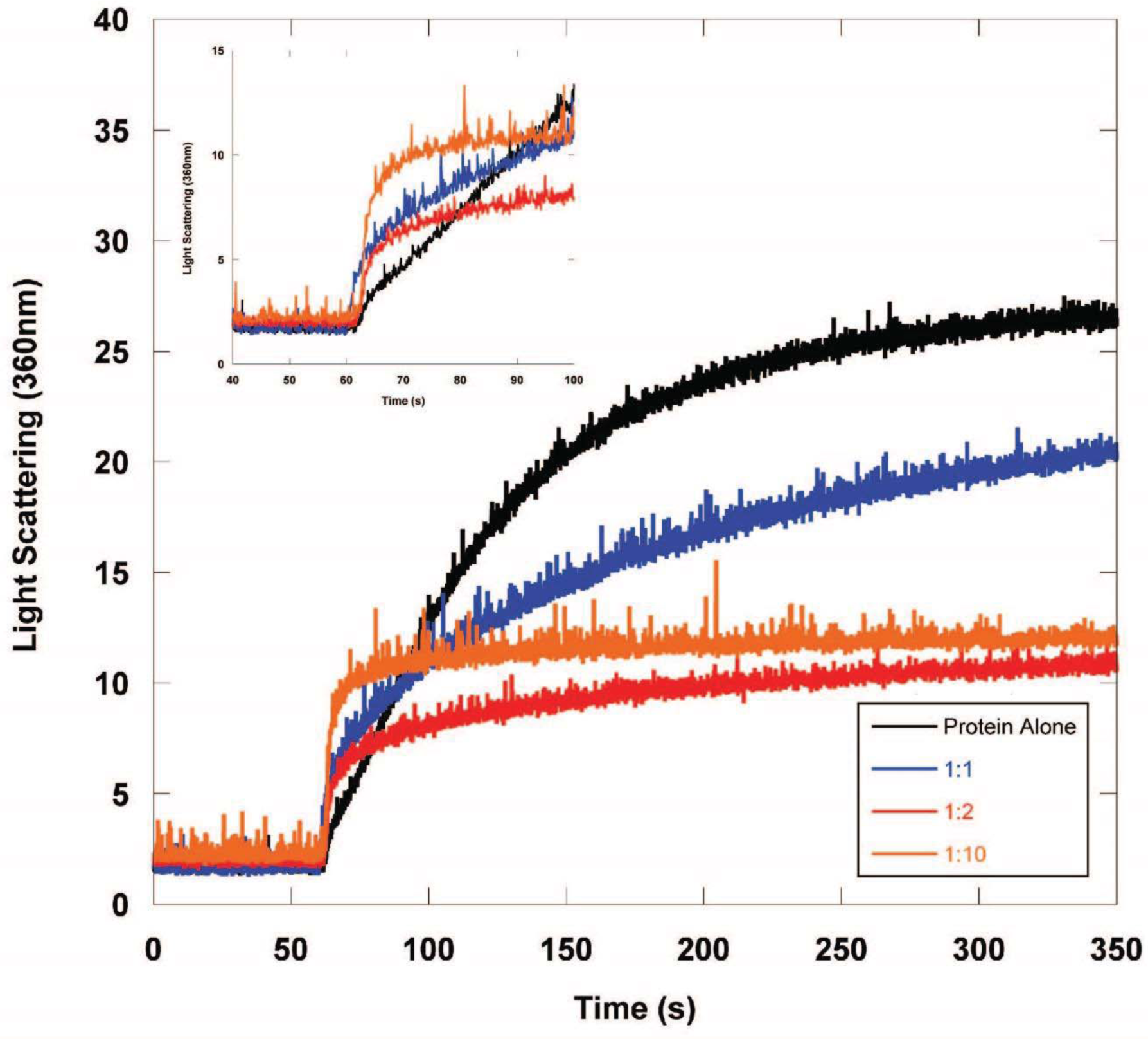
Chemical aggregation test of the concentration dependence of holdase activity of sequence 359, the best-performing holdase sequence. Concentration ratios are citrate synthase:DNA strand.

**Fig. S4.**
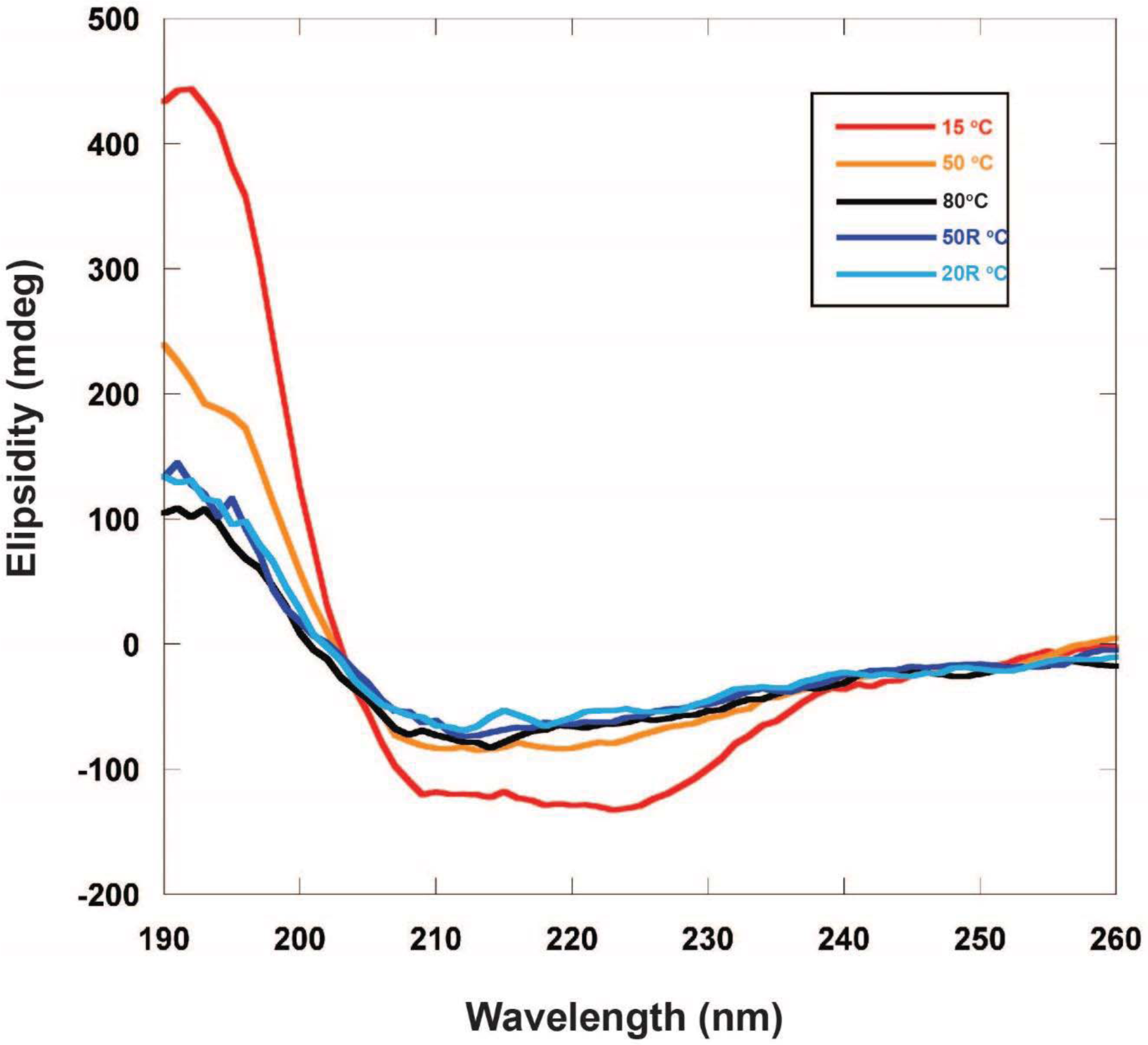
Circular dichroism of luciferase in the presence of quadruplex-forming sequence 359 throughout thermal denaturation. Luciferase retains partial beta sheet structure even at extreme temperatures, which is retained after returning to lower temperature.

**Fig. S5.**
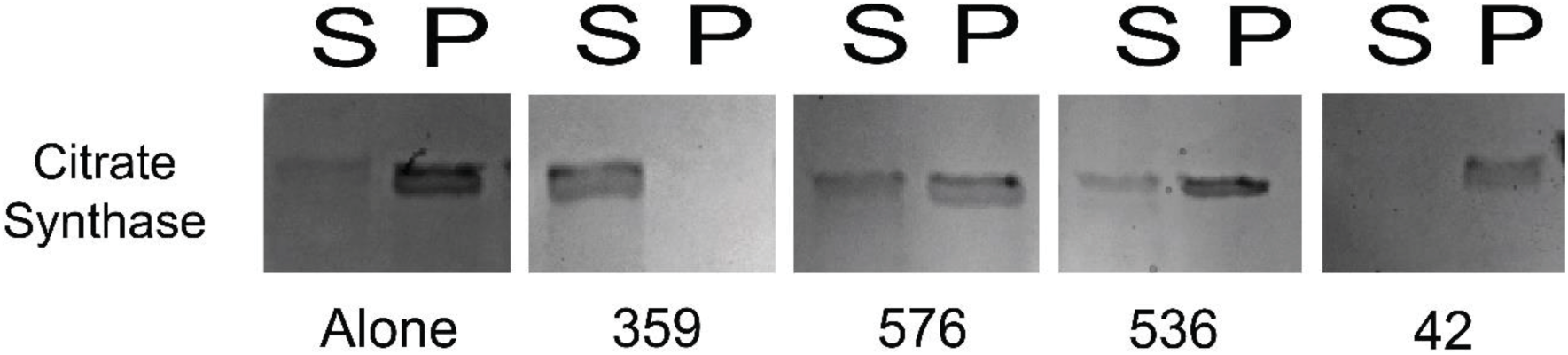
Prevention of protein aggregation in chemical spin down assay using citrate synthase. Sequences 359, 536, and 576 all displayed holdase activity and contain a polyG motif. Sequence 42 was used as a negative control, as it performed poorly as a holdase chaperone and did not contain a polyG motif.

